# Direct, bisulfite-free 5mC and 5hmC sequencing at single-cell resolution

**DOI:** 10.1101/2024.02.12.579864

**Authors:** Xiufei Chen, Jingfei Cheng, Linzhen Kong, Xiao Shu, Haiqi Xu, Masato Inoue, Marion Silvana Fernández Berrocal, Dagny Sanden Døskeland, Shivan Sivakumar, Yibin Liu, Jing Ye, Chun-Xiao Song

## Abstract

We report the first direct sequencing methodologies for quantitative detection of 5mC and 5hmC at single-base resolution and single-cell level, termed scTAPS (for 5mC + 5hmC) and scCAPS+ (for 5hmC specifically). With ∼90% mapping efficiency, our methods accurately benchmark 5mC and 5hmC profiles in CD8+ T and mES cells, respectively. Notably, scCAPS+ revealed a global increase in 5hmC within the hippocampus of aging mice, both in neurons and in non-neurons.

## Main

5-Methylcytosine (5mC) is a prevalent epigenetic modification in mammalian DNA, which is present in 70-80% of the symmetrical CpG dinucleotides^1^. 5-Hydroxymethylcytosine (5hmC) is a main derivative of 5mC by TET oxidation^2, 3^. Over the past two decades, extensive research has been dedicated to understanding the distinct roles played by 5mC and 5hmC in various biological processes^4, 5^. To charting 5mC and 5hmC at single-cell resolution, several sequencing methods based on bisulfite sequencing (BS-Seq) have been developed^6–8^. However, these methods encounter challenges such as substantial DNA damage and limited mapping efficiency. Recently, bisulfite-free methods^9, 10^ have emerged as alternatives to single-cell BS-Seq (scBS-Seq). It is worth noting that these methodologies still employ an indirect approach by converting unmodified cytosine (uC), resulting in low sequence complexity and high false positives.

Previously, we developed TET-assisted pyridine borane sequencing (TAPS)^11^ and chemical-assisted pyridine borane sequencing plus (CAPS+)^12, 13^ for 5mC and 5hmC detection, respectively, at bulk level. Here, by integrating Tn5 transposon-based fragmentation with TAPS and CAPS+, we present the first direct methodologies, namely single cell TAPS (scTAPS) and single cell CAPS+ (scCAPS+), for robust and precise profiling of DNA 5mC and 5hmC, respectively, at single-cell level in a bisulfite-free manner. Briefly, individual cells or nuclei are sorted via fluorescence-activated cell sorting (FACS), and then undergo lysis and fragmentation using a barcoded Tn5 transposons. Following gap-filling and DNA purification, barcoded DNA from 96 cells is pooled and TAPS or CAPS+ reaction is subsequently employed (Fig. 1a and Methods).

**Fig. 1:**
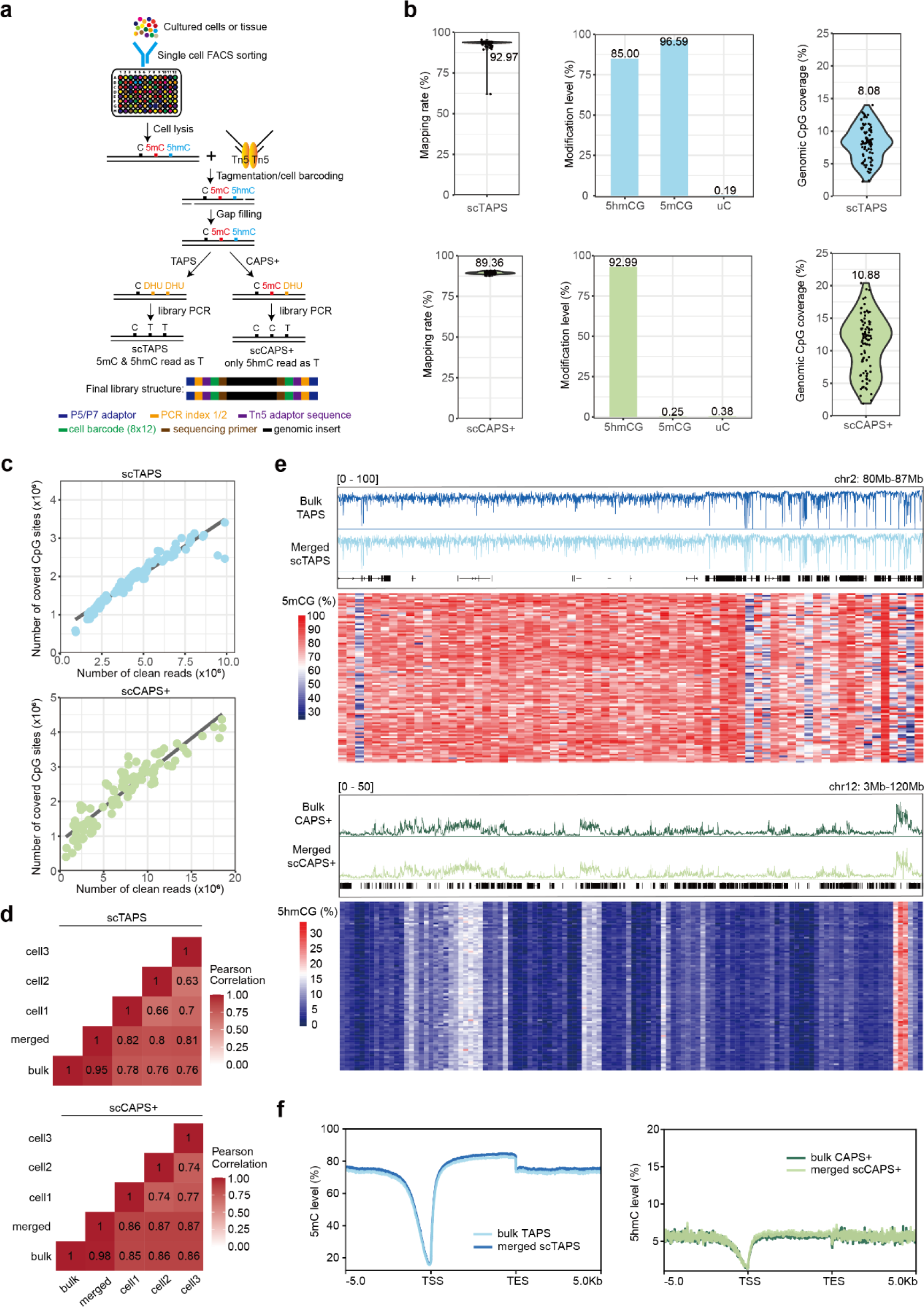
Validation of scTAPS and scCAPS+. **a.** Overview of the scTAPS and scCAPS+ methodologies. **b**. Violin plots showing the mapping rates (left) and genomic CpG coverage (right). Each point represents a single cell. Barplots (middle) showing conversion rates based on 5hmCG, 5mCG, and false positives on unmodified C spike-ins. The values above represent the means. Top: scTAPS. Bottom: scCAPS+. **c**, Saturation curve showing the number of covered CpG sites, and the number of cleaned reads sequenced in each cell. **d**, Heatmap showing Pearson correlation coefficient between bulk and single cell TAPS/CAPS+. **e**. 5mCG and 5hmCG level along chr2:80,000,000-87,000,000 in CD8+ T cell (top) and 5hmCG level along chr12: 3,000,000-120,000,000 in mES cells (bottom). IGV tracks in upper panel showing bulk 5mCG/5hmCG and merged 5mCG/5hmCG signals from scTAPS/scCAPS+. Heatmap in lower panels showing 5mCG/5hmCG levels in 100kb/1MB bins (column) across 96 CD8+ T/mES cells (row) in scTAPS/scCAPS+. Level is scaled by color. **f**. Metagene plots showing 5mCG and 5hmCG distribution along gene body from 5kb upstream of TSS (Transcription Start Sites) to 5kb downstream of TES (Transcription End Sites).

We first validated our methods in human peripheral blood CD8+ T cells for scTAPS and mouse embryonic stem cells (mES cells) for scCAPS+ (Fig. 1). Compared to previous methods^6–10^, we achieved the highest mapping efficiency in scTAPS (92.97%) and scCAPS+ (89.36%) (Fig. 1b). Notably, robust conversion rates were observed in both scTAPS (5mCG: 96.59%, 5hmCG: 85.00%) and scCAPS+ (5hmCG: 92.99%), along with very low false positive rates in scTAPS (uC: 0.19%) and scCAPS+ (uC: 0.38%, 5mCG: 0.25%) (Fig 1b). With a low sequencing depth of 4.8 and 7.7 million 120-base-pair (bp) paired-end reads per single cell, we achieved a mean coverage of 2.0 and 2.3 million CpG sites (8.08 % and 10.88% of the total CpG sites) in scTAPS and scCAPS+, respectively (Fig. 1b). We note that increased sequencing depth is likely to improve genomic coverage (Fig. 1c). Furthermore, 5mC/5hmC profiles in bulk TAPS and CAPS+ correlated strongly with merged single cells (Pearson’s r = 0.95) and 3 individual cell representatives (Fig 1d). The single cell 5mC/5hmC level closely aligned with those in the bulk samples (Fig 1e). Additionally, a concurrent reduction in both 5mC and 5hmC was observed in the proximity of transcription start site (TSS) regions (Fig. 1f), aligning with previous findings^11, 13^. Taken together, these findings illustrated scTAPS and scCAPS+ for accurately mapping 5mC and 5hmC in single cells.

The hippocampus is a critical brain region for learning and memory^14, 15^. To unravel aging related 5hmC dynamics in this region at the single-cell level, neurons and non-neurons were isolated from young (3 months) or aged (18 months) mice, followed by scCAPS+ analysis (Extended Data Fig. 1, Methods). We found hippocampus neurons exhibited higher 5hmC level compared to non-neurons (22.04% vs 9.29%) (Fig. 2a). Utilizing gene body 5hmC level, two distinct clusters (cluster 1 and cluster 2) were identified, closely corresponding to FACS-identified non-neuron and neuron cells (Fig. 2b). Next, we employed marker genes defined by 5hmC levels to annotate the two clusters using the Tabula Muris database^16^. It revealed that cluster 1 corresponds to Oligodendrocyte Precursor Cells (OPC) (CL: 0002453), while cluster 2 corresponds to neuronal cells (CL: 0000540) (Fig. 2c). Gene Ontology (GO) analysis revealed the enrichment of axon guidance and neuron projection guidance functions in both neuron and non-neuron marker genes (Fig. 2d, e), suggesting the significance of both cell types in maintaining fundamental neuronal functions. Non-neuron marker genes (*Cnksr3*, *Mob3b, Sema4d, Dock5*)^17^ exhibited significantly higher 5hmC levels in non-neurons, while neuron marker genes (*Cntnap2*, *Rbfox3, Syt1, Grm1*)^17^ displayed markedly elevated 5hmC levels in neurons (Fig. 2f, g, Extended Data Fig. 2c). Moreover, distinct separations between young and aged cells were observed in both non-neurons and neurons based on the 5hmC signal (Fig. 2h, i, Extended Data Fig. 2d). Correlation analysis revealed that genes exhibiting a positive correlation between expression and age^18^ (e.g., *Edil3* and *Prr5l*) tend to have higher levels of 5hmC in aged cells, and vice versa (e.g., *Epha3 and Srrm4*), particularly in non-neurons (Fig. 2j and Extended Data Fig. 2e).

**Fig. 2:**
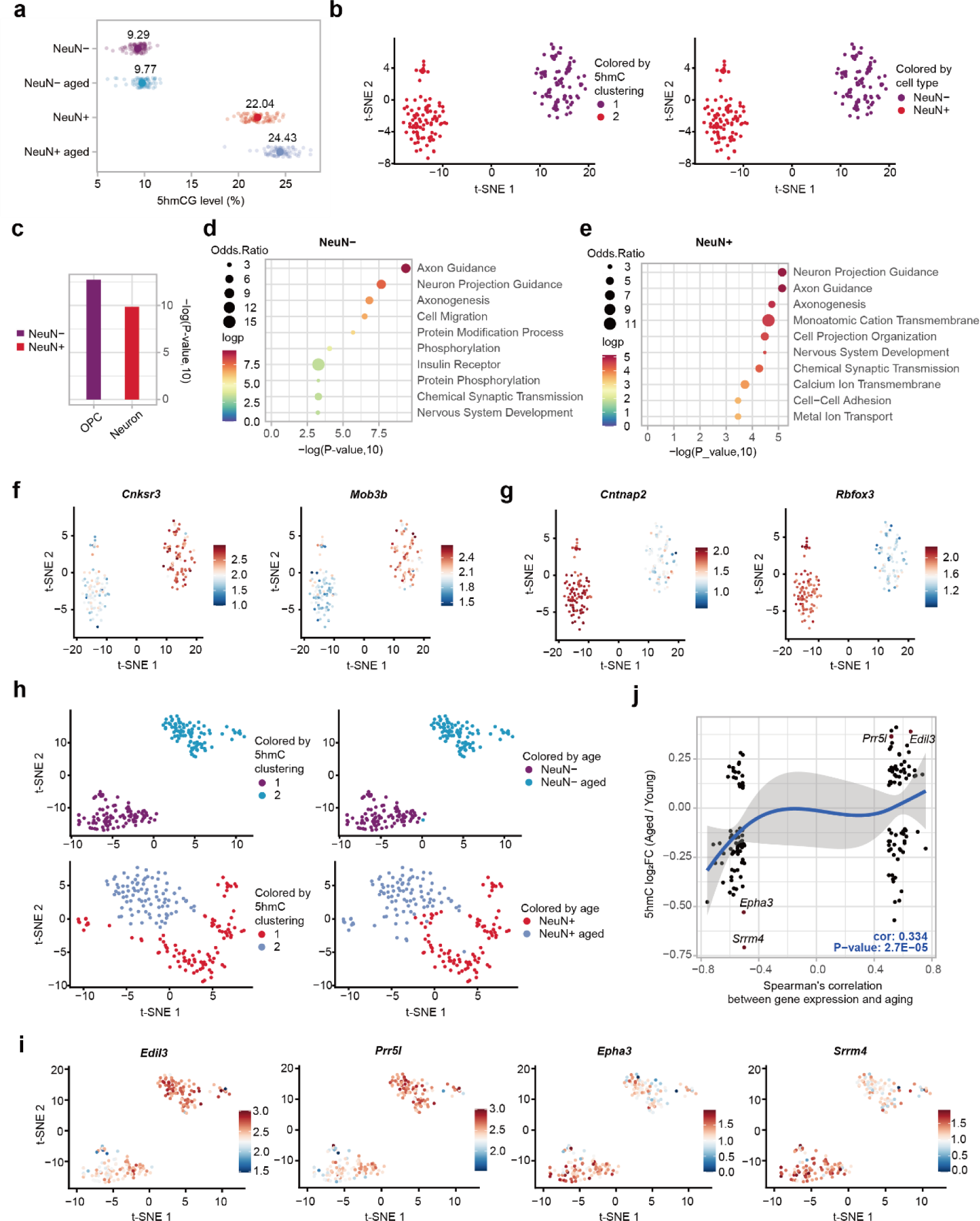
scCAPS+ unrevealed 5hmC dynamic in hippocampus during aging. **a.** Global 5hmC levels in young and aged non-neurons (NeuN-) or neurons (NeuN+). **b.** t-SNE plot showing the 5hmC clustering of young non-neurons and neurons. **c**. The top enriched cell ontology determined based on marker genes using the Tabula Muris dataset. OPC: Oligodendrocyte Progenitor Cell. **d, e**. Top 10 enriched GO terms for both non-neuron (d) and neuron (e) marker genes. **f, g**. t-SNE plot showing 5hmC levels of non-neurons marker genes (f) *Cnksr3* and *Mob3b* and neurons marker genes (g) *Cntnap2* and *Rbfox3*. **h**. t-SNE plot showing age related 5hmC clustering in both neuron (top) and non-neuron (bottom) cells. **i**. t-SNE plot showing 5hmC levels of *Edil3* and *Prr5l, Epha3 and Srrm4*, in young and aged non-neurons. **j**. The scatter plot showing the 5hmC changes in non-neurons during aging, along with the corresponding Spearman’s correlation coefficient depicting the relationship between gene expression and aging in the mouse brain. Pearson correlation coefficient and significance level were computed.

In summary, we present scTAPS and scCAPS+ as robust techniques for accurate and direct detection of5mC and 5hmC in single cells. By applying scCAPS+ to mouse hippocampus we revealed dynamics of 5hmC during aging. We anticipate that our methodologies will facilitate future investigations into the roles of 5mC and 5hmC in specific biological processes.

## Methods

### mES Cell culture

E14 mouse embryonic stem cells (mES cells) were cultured on gelatin-coated plates in DMEM supplemented with 15% FBS, 2 mM L-glutamine, 1% non-essential amino acids, 1% penicillin/streptavidin, 0.1 mM β-mercaptoethanol, 1,000 units/ml LIF (leukaemia inhibitory factor), 1 μM PD0325901, and 3 μM CHIR99021. The cells were maintained at 37°C and 5% CO_2_. Before cell sorting, the cells were harvested by centrifugation and passed through a 40 μm cell strainer (Falcon) to achieve single cell suspension.

### Animals

3 and 18 months old male mice with the C57BL/6N background were used in the study. Animals were housed with their littermates in 1717 × 545 × 2045 mm (LxWxH) cages with free food and water access in a dedicated room (temperature 22°C ± 1°C and humidity 55% ± 5%) with a 12h light/dark cycle (lights on 7 pm to 7 am). All animal experiments were conducted in accordance with the Norwegian Animal Welfare Act and approved by the Norwegian Animal Research Authority (FOTS 28340).

### Hippocampal tissue collection and nuclei isolation

Mice were anesthetized using isoflurane (Baxter, Oslo, Noway) and subsequently killed by an intraperitoneal overdose of pentobarbital (>200mg/kg body weight). The mouse brain was directly extracted without intracardial perfusion, and the hippocampal regions were micro-dissected in cold PBS. Nuclei from freshly dissected hippocampus (HPC) were isolated with EZ PREP kit (Sigma, Cat #NUC-101). Briefly, two single-side HPC from two mice at 3m or 18m age were resuspended in 1 ml ice-cold EZ lysis buffer, homogenized using a glass Dounce tissue grinder (15 times with the loose pestle and 15 times with the tight pestle), and incubated on ice for 5 min. Homogenate was strained through a 30μm cell strainer (Miltenyi Biotech) and centrifuged at 850xg for 10 minutes (4°C) to pellet nuclei. Nuclei were washed in 1ml ice-cold EZ lysis buffer with a 5-min incubation on ice, and then pelleted by centrifugation (850xg, 10min, 4°C).

### Nuclei staining and Flow cytometry-based cell sorting (FACS)

Nuclei were resuspended in 1ml staining buffer (1x PBS supplemented with 1% nuclease-free BSA) and incubated at 4°C for 10 min (blocking). Mouse anti-NeuN antibody (Merck Millipore, MAB377, Clone A60, RRID: AB_2298772) was added to the nuclei at a final dilution of 1:500, and nuclei suspensions were incubated at 4°C for 30 minutes. Nuclei were pelleted with centrifugation (450xg, 10min, 4°C) and washed once in the staining buffer. After washing and pelleting, nuclei were resuspended in 1ml staining buffer containing the secondary antibody (goat anti-mouse IgG1, Alexa Fluor™ 488, ThermoFisher Scientific, RRID: AB_2535764) at a final dilution of 1:1000. After a 30-min incubation at 4°C, nuclei were again pelleted (450xg, 10min, 4°C) and washed once in the staining buffer. Prior to FACS, nuclei were resuspended in 1ml staining buffer containing DAPI at a final concentration of 0.1ug/ml and filtered through a 30μm cell strainer. Single nuclei were captured by gating on DAPI-positive events, and then gating on Alexa Fluor 488 (NeuN) signal. NeuN+ and NeuN-nuclei were sorted in Optical 96-Well plates (ThermoFisher Scientific) containing 4µl of lysis buffer consisting of 16.67mM TAPS-NaOH buffer (pH 8.5, Alfa Aesar, Cat: J63268), 8.33mM MgCl_2_(Invitrogen), 0.67x NEB buffer 4 (NEB, B7004S), 0.13% Triton X-100 (sigma, X100-500ML), and 1 μg Qiagen protease in each well.

### The preparation of Spike-ins and 2kb filler DNA for scTAPS and scCAPS+

Fully methylated lambda phage DNA, 2 kb unmodified DNA and 2kb filler DNA were made as previously described^11, 13^. To make 5hmC spike-in, three DNA oligos (oligo1, 5’-GTCTCGTGGGCTCGGCTGTCCCTGTCCCTATCGCTCACCGTCTCCGCCTCAGATGTGTATAAGAGACAGCATGCCAGCGTGCGCC; oligo2, 5’-TGAGTGTTTCATCCTCTGTCTCTTATACACATCTGCCGTCCTCGATCGCGTCAGTACGCGTGGAGACGCTGCCGACGA; oligo3: 5’-AGGATGAAACACTCAGGCTAGTGAGAATGAAGGGATATGTTTGTAAGATGGTCNNGNATCTTGGGTTGTGTGGTGGATGTTGGCGTTGGTGGGTTTCAGAGTTGGGGCGCACGCTGGCATG; 5 μM each) from IDT were annealed in 1x New England Biolabs (NEB) buffer 2 (20 μl in total). Subsequently, the annealed oligos were incubated with 10 mM 5-hydroxymethyl-2’-dCTP (Zymo Research), along with dGTP, dATP, dTTP (NEB), and 10 U Klenow exo^−^ (M0212L) in a 50 μl reaction at 37 °C for 1 h. T4 ligation was then carried out at room temperature for 30 mins. Following purification, the DNA underwent End Repair and A-Tailing (ER&T), followed by ligation with Tn5 i7/i5 adaptors sequence with T in the 3’ end. The resulting spike-in was finally purified using 1.8× AMPure XP beads, following the provided guidelines. the final 5hmC spike in sequence: GTCTCGTGGGCTCGGCTGTCCCTGTCCCTATCGCTCACCGTCTCCGCCTCAGATGTGTATAAGAGACAGCATGCCAGCGTGCGCCCCAACTCTGAAACCCACC AACGCCAACATCCACCACACAACCCAAGATNhmCNNGACCATCTTACAAA CATATCCCTTCATTCTCACTAGCCTGAGTGTTTCATCCTCTGTCTCTTATACACATCTGCCGTCCTCGATCGCGTCAGTACGCGTGGAGACGCTGCCGACGA.

### Tn5 protein purification

Tn5 protein is purified as previously described with some modifications^19^, NEB C3013 cells were transformed with the pTBX1-Tn5 plasmid, Tn5 protein is induced by adding 250 µl of 1 M IPTG ((Isopropyl β-D-1-thiogalactopyranoside) into 1 L LB culture when the A600 reached 0.7-0.9, incubation for 16-18 hours at 16°C in a shaking incubator. The cells were pelleted at 4000 rpm for 30 min and resuspended in 50 mL of cold HEGX buffer (20mM HEPES-KOH pH 7.2, 0.8M NaCl, 1mM EDTA, 10% glycerol, 0.2% Triton X-100, 1× complete Protease Inhibitor Cocktail). Cell lysate was sonicated with a BioRuptor sonicator for 20 mins at 40% Amp, 5 sec on/10 sec off (on ice), and then centrifuged at 35,000*g* for 60 min at 4°C. The supernatant was collected and precipitated with 2.1 mL of 10% neutralized PEI in a dropwise manner. The resulting mixture was centrifuged at 21,500*g* for 10 min at 4°C, and the supernatant was pre-incubated with 10 mL of washed chitin beads for 3 hours in a cold room. The sample was loaded onto a gravity column, washed with 500 mL of cold HEGX buffer, and equilibrated with 5 mL of HEGX with 100 mM DTT (Dithiothreitol). The column was incubated for Tn5 elution at 4°C for 48 hours. The eluted sample was dialyzed overnight at 4°C in 2 L of 2x Tn5 dialysis buffer (100mM HEPES-KOH pH 7.2, 0.2M NaCl, 0.1mM EDTA, 20% glycerol, 2mM fresh added DTT) using a Spectra/Por 6-8kD dialysis membrane. The dialyzed sample was quantified for concentration using Nanodrop, mixed with an equal volume of 100% glycerol for homogeneity, frozen in liquid nitrogen, and stored in aliquots at −80 °C before use.

### Tn5 transposome assembly

96 combinatory Tn5 assembly were performed as described before^19^. and their sequences are presented herein. Briefly, 8×12 Tn5MEDS-i5/i7 and Tn5MEDS-Rev oligos were purchased from IDT, and annealing was conducted in 1X TE buffer with a 25 μl total volume (95°C for 5 minutes, followed by a gradual decrease in temperature at a rate of 1°C per 12 seconds for 180 cycles, and cooling down to 4°C). The Tn5 assembly took place in a 96-well plate, combining 2 μL of Tn5 (10 ng/μL), 2 μL of pre-annealed Tn5MEDS-i5 and i7 oligos, 25 μL of glycerol, and 29 μL of 2x Tn5 dialysis buffer to reach a total volume of 60 μL. This reaction was incubated at room temperature for 1 hour, followed by a 7-fold dilution to produce 1x preassembled Tn5 complexes. Preassembled Tn5 complexes were stored at −80 °C and demonstrated stability over a one-month period. 1 μL of 1x preassembled Tn5 was always added to each well for scTAPS and scCAPS+ experiment,

### 8×12 Tn5 i5(1-8)/i7(1-12) sequence (5’-3’)^20^

P5_i5_1_Universal_Connector_A_C15_ME,

/5Phos/TCGTCGGCAGCGTCTCCACGCTATAGCCTGCGATCGAGGACGGCAG ATGTGTATAAGAGACAG

P5_i5_2_Universal_Connector_A_C15_ME,

/5Phos/TCGTCGGCAGCGTCTCCACGCATAGAGGCGCGATCGAGGACGGCAG ATGTGTATAAGAGACAG

P5_i5_3_Universal_Connector_A_C15_ME,

/5Phos/TCGTCGGCAGCGTCTCCACGCCCTATCCTGCGATCGAGGACGGCAG ATGTGTATAAGAGACAG

P5_i5_4_Universal_Connector_A_C15_ME,

/5Phos/TCGTCGGCAGCGTCTCCACGCGGCTCTGAGCGATCGAGGACGGCAG ATGTGTATAAGAGACAG

P5_i5_5_Universal_Connector_A_C15_ME,

/5Phos/TCGTCGGCAGCGTCTCCACGCAGGCGAAGGCGATCGAGGACGGCA GATGTGTATAAGAGACAG

P5_i5_6_Universal_Connector_A_C15_ME,

/5Phos/TCGTCGGCAGCGTCTCCACGCTAATCTTAGCGATCGAGGACGGCAG ATGTGTATAAGAGACAG

P5_i5_7_Universal_Connector_A_C15_ME,

/5Phos/TCGTCGGCAGCGTCTCCACGCCAGGACGTGCGATCGAGGACGGCAG ATGTGTATAAGAGACAG

P5_i5_8_Universal_Connector_A_C15_ME,

/5Phos/TCGTCGGCAGCGTCTCCACGCGTACTGACGCGATCGAGGACGGCAG ATGTGTATAAGAGACAG

P7_i7_1_Universal_Connector_B_D15_ME,

/5Phos/GTCTCGTGGGCTCGGCTGTCCCTGTCCCGAGTAATCACCGTCTCCGCCTCAGATGTGTATAAGAGACAG

P7_i7_2_Universal_Connector_B_D15_ME,

/5Phos/GTCTCGTGGGCTCGGCTGTCCCTGTCCTCTCCGGACACCGTCTCCGCCTCAGATGTGTATAAGAGACAG

P7_i7_3_Universal_Connector_B_D15_ME,

/5Phos/GTCTCGTGGGCTCGGCTGTCCCTGTCCAATGAGCGCACCGTCTCCGCCTCAGATGTGTATAAGAGACAG

P7_i7_4_Universal_Connector_B_D15_ME,

/5Phos/GTCTCGTGGGCTCGGCTGTCCCTGTCCGGAATCTCCACCGTCTCCGCCTCAGATGTGTATAAGAGACAG

P7_i7_5_Universal_Connector_B_D15_ME,

/5Phos/GTCTCGTGGGCTCGGCTGTCCCTGTCCTTCTGAATCACCGTCTCCGCCTCAGATGTGTATAAGAGACAG

P7_i7_6_Universal_Connector_B_D15_ME,

/5Phos/GTCTCGTGGGCTCGGCTGTCCCTGTCCACGAATTCCACCGTCTCCGCCTCAGATGTGTATAAGAGACAG

P7_i7_7_Universal_Connector_B_D15_ME,

/5Phos/GTCTCGTGGGCTCGGCTGTCCCTGTCCAGCTTCAGCACCGTCTCCGCCTCAGATGTGTATAAGAGACAG

P7_i7_8_Universal_Connector_B_D15_ME,

/5Phos/GTCTCGTGGGCTCGGCTGTCCCTGTCCGCGCATTACACCGTCTCCGCCTCAGATGTGTATAAGAGACAG

P7_i7_9_Universal_Connector_B_D15_ME,

/5Phos/GTCTCGTGGGCTCGGCTGTCCCTGTCCCATAGCCGCACCGTCTCCGCCTCAGATGTGTATAAGAGACAG

P7_i7_10_Universal_Connector_B_D15_ME,

/5Phos/GTCTCGTGGGCTCGGCTGTCCCTGTCCTTCGCGGACACCGTCTCCGCCTCAGATGTGTATAAGAGACAG

P7_i7_11_Universal_Connector_B_D15_ME,

/5Phos/GTCTCGTGGGCTCGGCTGTCCCTGTCCGCGCGAGACACCGTCTCCGCCTCAGATGTGTATAAGAGACAG

P7_i7_12_Universal_Connector_B_D15_ME,

/5Phos/GTCTCGTGGGCTCGGCTGTCCCTGTCCCTATCGCTCACCGTCTCCGCCTCAGATGTGTATAAGAGACAG pMENTS,

/5Phos/CTGTCTCTTATACACATCT

### mTET1 protein expression

The mTET1 purification is conducted as previously described^11^. Briefly, the mTet1CD catalytic domain (NM_001253857.2, 4371–6392) was integrated into the pcDNA3-Flag vector using KpnI and BamH1 restriction sites. Subsequently, 1mg of plasmid was transfected into 1L of Expi293F cell culture at a density of 1×10^6^ cells ml^−1^. The cells were cultured for 48 h at 37 °C, 170 r.p.m., and 5% CO2. Following incubation, cells were harvested by centrifugation, resuspended in a lysis buffer (50 mM Tris–Cl pH 7.5, 500 mM NaCl, 1× complete Protease Inhibitor Cocktail, 1 mM PMSF, 1% Triton X-100), and incubated on ice for 20 min. Next, the lysed cell solution was clarified by centrifugation at 30,000*g* and 4 °C for 30 min. The resulting supernatant was purified using ANTI-FLAG M2 Affinity Gel. Pure protein was eluted in the buffer (20 mM HEPES pH 8.0, 150 mM NaCl, 0.1 mg ml^−1^ 3× Flag peptide, 1× complete Protease Inhibitor Cocktail, and 1 mM PMSF). The collected fractions were then concentrated, and buffer exchanged into a final buffer of 20 mM HEPES pH 8.0, 150 mM NaCl, and 1 mM DTT. The concentrated protein was mixed with glycerol (30% v/v), frozen in liquid nitrogen, and stored in aliquots at −80 °C.

### scTAPS and scCAPS+

For scTAPS, 96 cells were pooled for DNA purification and subjected to mTET1 oxidation as previously described^11^. A reaction mixture of 50 μl was prepared, comprising 50 mM HEPES buffer (pH 8.0), 100 μM ammonium iron (II) sulfate, 1 mM α-ketoglutarate, 2 mM L-ascorbic acid, 2.5 mM DTT, 1.2 mM ATP, 100 mM NaCl, and 4 μM mTet1CD. The reaction was conducted at 37°C for 80 minutes. Subsequently, 2.0 U of Proteinase K (New England Biolabs) was introduced to the reaction mixture and incubated at 50°C for 1 hour to stop the oxidation process. The oxidized DNA was purified using 1.8x AMPure XP beads, following the provided guidelines. To ensure complete oxidation, a second round of oxidation was performed. The double-oxidized DNA was purified and then underwent borane reduction and purification using a Zymo column with Oligo Binding Buffer.

For scCAPS+, 96 cells were pooled for DNA purification and subjected to a series of chemical reactions to convert 5hmC into 5fC and then to 5caC as previously described with some modifications^13^. DNA were incubated with 4-acetamido-2,2,6,6-tetramethylpiperidine-1-oxoammonium tetra-fluoroborate (ACT+ BF_4_-) in a 25 μl solution comprising sodium phosphate buffer at 25°C for 16 hours. The resultant oxidized DNA was purified using 1.8× AMPure XP Beads. Then Pinnick oxidation is conducted in a 30 μl reaction mixture containing sodium acetate buffer, NaClO_2_, and 2-methyl-2-butene at 25°C for 16 hours. The purified product then underwent borane reduction and purification using a Zymo column with Oligo Binding Buffer.

### scTAPS and scCAP+ library construction and sequencing

Individual cells were isolated into a 96-well plate through FACS as mentioned above. Following centrifugation at 2,000 rpm for 5 minutes, the plates were either preserved at -80°C or progressed to subsequent procedures. Single cell was lysate by incubating at 50°C for 3 h, followed by incubation at 75°C for 30 min, 80°C for 15 min, and a 4°C hold. Tagmentation was performed at 50°C for 15 minutes in a reaction mix consisting of 1x TAPS buffer (10mM TAPS-NaOH, 5mM MgCl_2_), 6% PEG (polyethylene glycol) 3,350, 1ul 1x preassembled Tn5, and H_2_O to a total volume of 10 μ. Tn5 was stripped off by adding 1.11 μL of 1.0% SDS (Sodium Dodecyl Sulfate), followed by incubating at room temperature for 5 minutes. Neutralization was performed by adding 1.24 μL of 5.0% NP40 or Triton X-100, followed by incubating at room temperature for 5 minutes. Subsequently, 12.35 μL of 2x Phusion High-Fidelity PCR Master Mix with HF Buffer (Thermo Scientific, F531L) was added, followed by incubating at 72°C for 5 minutes, and a 4°C hold. The 96 reactions were combined into a 50-mL BD tube and supplemented with 2-kb filler DNA (200 ng) and 0.1% spike in DNAs. Following purification, the TAPS or CAPS+ reaction, was employed as mentioned, and the final sequencing library was amplified utilizing the KAPA HiFi HotStart Uracil+ ReadyMix PCR Kit along with index primers from the Nextera XT Index Kit (Illumina, FC-131-1001). The amplification protocol involved an initial step at 98°C for 45 seconds, followed by 13 cycles (98°C for 10 seconds, 60°C for 15 seconds, and 72°C for 1 minute), then incubation at 72°C for 3 minutes, 4°C hold. The library was subjected to purification and size selected to 500-700bp by 0.5x-0.25x Ampure XP beads. Custom sequencing primers are used for custom Nova Sequencing (Read 1 Sequencing Primer: GCGATCGAGGACGGCAGATGTGTATAAGAGACAG; Read 2 Sequencing Primer: CACCGTCTCCGCCTCAGATGTGTATAAGAGACAG; Index 1 Sequencing Primer: CTGTCTCTTATACACATCTGAGGCGGAGACGGTG; Index 2 sequencing primer: CTGTCTCTTATACACATCTGCCGTCCTCGATCGC).

### Data analysis for scTAPS and scCAPS+

Pre-processing: Raw sequenced reads were demultiplexed using demuxbyname2.sh script from BBMap (version 38.50; https://www.osti.gov/biblio/1241166). Demultiplexed reads were processed with fastp^21^ package using default parameters to remove low-quality bases. The trimmed reads were aligned to reference genome using bwa^22^ with default parameters. Reference sequence for mouse genome were downloaded from https://hgdownload.cse.ucsc.edu/goldenpath/mm9/bigZips/; reference sequence for human CD8+ T-cell were downloaded from: ftp://ftp.ncbi.nlm.nih.gov/genomes/all/GCA/000/001/405/GCA_000001405.15_GRC h38/seqs_for_alignment_pipelines.ucsc_ids/GCA_000001405.15_GRCh38_no_alt_an alysis_set.fna.gz. Uniquely mapped reads were filtered using samtools^23^ tools with MAPS>=10. PCR duplicates were called using Picard (2.23.0-Java-11) Mark Duplicates (https://broadinstitute.github.io/picard/). Methylation was called using a custom R script used in our previous project^24^. 10bp in start or end of reads were excluded for methylation calling. Snakemake^25^ pipeline file with detailed steps is provided in the code availability section.

Clustering for mouse neuron scCAPS+: Cells with an insufficient number of properly mapped reads (<500,000) or an excessively high count (>3,000,000) were systematically excluded from downstream analysis. The 5hmCG level for each annotated protein-coding genes were computed by dividing the sum of methylated base calls by the total base calls across gene body. Gene annotation was downloaded from https://ftp.ebi.ac.uk/pub/databases/gencode/Gencode_human/release_43/gencode.v43. basic.annotation.gtf.gz. Genes with total calls less than 5 were excluded for analysis. The Seurat^26^ package was used for downstream analysis with parameters specified in the code availability section for reproducibility-related details. Marker genes were identified by the FindMarkers function using the min.pct=0.25 parameter. For subsequent functional enrichment analysis, genes with an adjusted p-value (p_val_adj) less than 0.05 were selected. The enrichment analysis was conducted using the enrichR package^27^ against the GO_biological_Process_2023 ontology and the Tabula_Muris database (https://tabula-muris.ds.czbiohub.org/). The age-correlated genes and the correlation between their expression and age were downloaded from a previous study^18^. Annotation: Annotated genes were downloaded from RefSeq database (https://www.ncbi.nlm.nih.gov/refseq/MANE/) for mESCs and Gencode database (https://ftp.ebi.ac.uk/pub/databases/gencode/Gencode_human/release_43/gencode.v43.basic.annotation.gtf.gz) for CD8+ T cells. 5mCG/5hmCG scores within gene bodies, 5 kb upstream of TSS and 5 kb downstream regions of transcription termination sites (TTS) sites were calculated by computeMatrix in deepTools (v.3.3.1)^28^ with parameters (--beforeStartLength 5000 --regionBodyLength 5000 --afterRegionStartLength 5000 --binSize 10). The plots were created with plotProfile function. Chromatin states for mES and CD8+ cells were downloaded from https://github.com/gireeshkbogu/chromatin_states_chromHMM_mm9^29^, and https://github.com/ernstlab/full_stack_ChromHMM_annotations^30^, respectively, Aggregated 5mC/5hmC signals were calculated in each chromatin state in each cell.

## Acknowledgements

We would like to acknowledge B.M. for helping with single cell sorting. Our work is supported by Ludwig Institute for Cancer Research, Cancer Research UK (C63763/A26394), and National Institute for Health Research (NIHR) Oxford Biomedical Research Centre (BRC). H.X. and L.K. are supported by China Scholarship Council (CSC). M.I. is supported by the Nakajima foundation. Y.L. is supported by National science Foundation of China (No. 22307099).

## Author Contribution

X.C., Y.L., J.Y. and C.S. conceived the study and designed the experiment. X.C. carried out most experiments with help from H.X., M.I., M.S.F.B., D.S.D., S.S. L.Z and J.F. developed the algorithm and analyzed data. X.C. and X.S. wrote the manuscript.

## Figure legends

**Extended Data Fig. 1.**
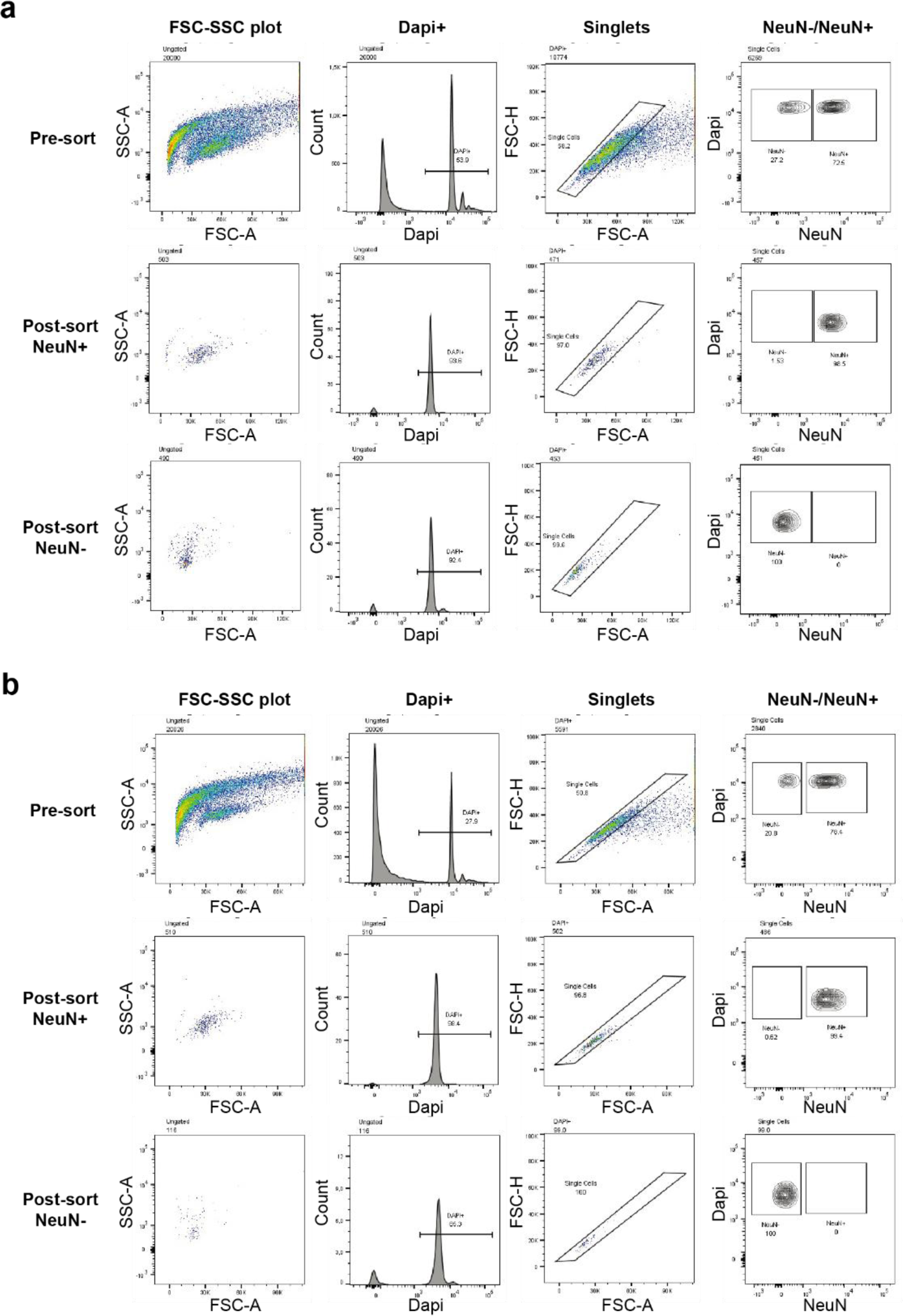
Separation of neurons (NeuN+) and non-neurons (NeuN-) by FACS. Nuclei isolated from hippocampal tissues of 3m (a) and 18m (b) mice were stained with the anti-NeuN antibody and sorted by FACS. Single nuclei were captured by gating on DAPI-positive events, and then gating on Alexa Fluor 488 (NeuN) signal. Plots showing the pre-sorted and post-sorted NeuN+/NeuN-nuclei were listed.

**Extended Data Fig. 2:**
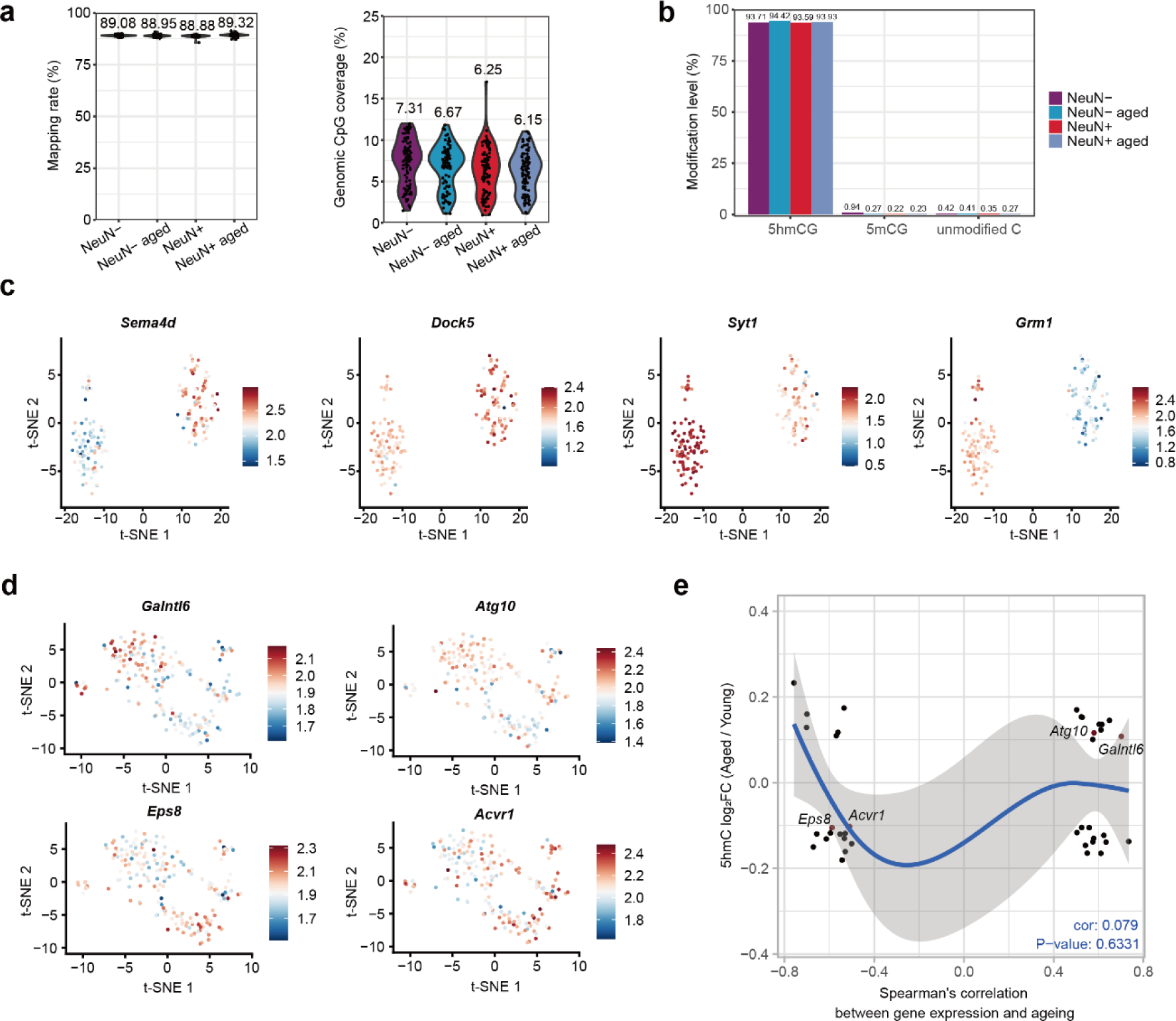
scCAPS+ unrevealed 5hmC dynamic in hippocampus during aging. **a.** Violin plots showing the mapping rates (left) and genomic CpG coverage (right) of scCAPS+ in mouse hippocampus, each point represents a single cell. The means are shown above. **b**. Conversion rate and false positive of scCAPS+ in mouse hippocampus. **c.** t-SNE plot showing 5hmC levels of non-neurons marker genes *Sema4d* and *Dock5* and neurons marker genes *Syt1* and *Grm1*. **d**. t-SNE plot showing 5hmC levels of *Galntl6 and Atg10*, *Eps8* and *Acvr1* in young and aged neurons. **e**. Scatter plot showing the 5hmC changes in neuronal cells during aging, along with the corresponding Spearman’s correlation coefficient depicting the relationship between gene expression and aging in the mouse brain. Pearson correlation coefficient and significance level were computed.

